# Spatial autocorrelation of the environment influences the patterns and genetics of local adaptation

**DOI:** 10.1101/2023.05.29.542754

**Authors:** Tom R. Booker

## Abstract

Environmental heterogeneity can lead to spatially varying selection, which can, in turn, lead to local adaptation. Population genetic models have shown that the pattern of environmental variation in space can strongly influence the evolution of local adaptation. In particular, when environmental variation is highly autocorrelated in space local adaptation will more readily evolve. Despite this long-held prediction, the evolutionary genetic consequences of different patterns of environmental variation have not been thoroughly explored. In this study, simulations are used to model local adaptation to different patterns of environmental variation. The simulations confirm that local adaptation is expected to increase with the degree of spatial autocorrelation in the selective environment, but also show that highly heterogeneous environments are more likely to exhibit high variation in local adaptation, a result not previously described. Spatial autocorrelation in the environment also influences the evolution and genetic architecture of local adaptation, with different combinations of allele frequency and effect size arising under different patterns of environmental variation. These differences influence the ability to characterise the genetic basis of local adaptation in different environments. Finally, I analyse a large-scale provenance trial conducted on lodgepole pine and find suggestive evidence that spatially autocorrelated environmental variation leads to stronger local adaptation in natural populations of lodgepole pine. Overall, this work emphasizes the profound importance that the spatial pattern of selection can have on the evolution of local adaptation and how spatial autocorrelation should be considered when formulating hypotheses in ecological and genetic studies.

**Lay Summary:** Many species exhibit local adaptation to environmental variation across their ranges. Theoretical population genetics predicts that the evolution of local adaptation and patterns of genetic variation underlying it will be influenced by the spatial pattern of variation across a species’ range. However, this prediction has not been thoroughly explored for cases of complex heterogeneous landscapes. In this paper, I analyse simulations and empirical data to characterise the effects that the spatial pattern of environmental variation can have on the evolution of local adaptation and the genetics underlying it. From these analyses, I show that the pattern of environmental variation influences the average level of local adaptation, variation in local adaptation as well as the genetics underlying this important phenomenon.

## Introduction

Local adaptation is an important phenomenon in the natural world. Along with phenotypic plasticity, local adaptation can dictate the extent of environmental heterogeneity that a species can tolerate, shape its geographic range (Kirkpatrick & Barton, 1997), and help predict how it will respond to changing environments (Rellstab et al., 2021). In forest trees, for example, local adaptation is widely observed (Leites & Benito Garzón, 2023) and is central to plans for adapting forestry practice in light of climate change (O’Neill & Gómez-Pineda, 2021; Ying & Yanchuk, 2006). Local adaptation can be defined as a kind of genotype-by-environment interaction for fitness, where individuals have higher chances of survival and/or reproduction when they are reared at home as opposed to away, though several other definitions are used in the literature (Blanquart et al., 2013; Kawecki & Ebert, 2004). Local adaptation is a property of a particular population at a particular point in time rather than a property of a species as a whole. For example, populations at range edges are often expected to be maladapted to their conditions, while populations in the core of a range may be well adapted (Angert et al., 2020). Locally adapted populations may harbour genetic variation that could help buffer susceptible ones against the detrimental effects of climate change (Aitken & Whitlock, 2013), which are already wreaking havoc on important species around the world (Hartmann et al., 2022). A deep understanding of local adaptation, the agents that have given rise to it and the genetics that underpin this phenomenon is thus important for our understanding of biodiversity and for species management and conservation in the Anthropocene (Aitken & Whitlock, 2013; Exposito-Alonso, 2023; Wadgymar et al., 2022).

The ultimate cause of local adaptation is variation in the environment. Whether it is biotic (e.g. disease/parasite prevalence or intraspecific competition) or abiotic (e.g. climate, geology or photoperiod), variation in the environment may lead to spatially varying selection pressures where phenotypic optima differ over a landscape. Such variation in selection across space has been well documented (e.g. Siepielski et al., 2013) and there are, of course, myriad aspects of the environment that could conceivably induce spatially varying selection, many of which would be highly inter-correlated. However, while there is an infinite number of ways to describe the environment, most of these may be functionally disconnected from a species’ biology. A recent review by Wadgymer et al (2022) highlighted a critical gap in our knowledge of local adaptation - that the aspects of environmental variation that have given rise to local adaptation (what they term the ‘agents of selection’) are unknown in most cases. For example, in the absence of experimental evidence many genetic studies have assumed that various climatic measures recorded in databases such as WorldClim correspond to agents of selection and search for the genetic basis of local adaptation using those data (Lasky et al., 2023). However, that a particular aspect of environmental variation could conceivably induce spatially varying selection is not a guarantee that it will have led to local adaptation.

Population genetic studies have revealed numerous factors that can influence the evolution of local adaptation in a particular location, the most important being the strength of selection and rates of gene flow. The strength of natural selection is of foremost importance because larger fitness consequences for deviating from the optimal phenotype in a particular location can potentially lead to greater evolutionary change (Falconer & MacKay, 1995). The rate of gene flow is important because migration into a region experiencing idiosyncratic selection can overwhelm that selection, preventing regional trait differences (i.e. local adaptation) from accumulating (Nagylaki, 1975; Slatkin, 1978; Wright, 1931; Yeaman, 2015). In discrete space models, the evolution of local adaptation in a particular location depends on the ratio of gene flow from dissimilar environments (*m*) to the strength of selection (*s*), *m/s* (Slatkin, 1978; Wright, 1931; Yeaman, 2015). In models of continuous space, patterns of dispersal relative to the strength of selection can be used to determine the minimum size a region experiencing idiosyncratic selection needs to be for locally adaptive differences to accumulate, the so-called “characteristic length” of a cline. Specifically, when dispersal is modelled as a diffusion process, the characteristic length is the standard deviation of dispersal distances (σ) relative to the square root of the strength of selection (i.e. 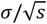). Such characteristic lengths can be described for individual alleles (Nagylaki, 1975; Slatkin, 1973) or for stabilising selection acting on polygenic traits (Barton, 1999; Slatkin, 1978), see reviews by Felsenstein, (1976) and Lenormand (2002). Furthermore, the mean values of polygenic traits will more closely track changes in phenotypic optima over space if those changes are small (Barton, 1999; Slatkin, 1978). Because natural populations often inhabit large spatial ranges encompassing complex patterns of environmental variation, relative rates of gene flow among regions of high environmental similarity or dissimilarity will vary across the landscape, influencing the evolution of local adaptation.

That the spatial pattern of environmental heterogeneity will influence the evolution of local adaptation has been recognised since at least the 1960s (Antonovics, 1971; Antonovics & Bradshaw, 1970; Forester et al., 2016; Hadfield, 2016; Levins, 1966; Schiffers et al., 2014). Indeed, the overall level of local adaptation a species exhibits can be strongly affected by the pattern of environmental variation over space (Forester et al., 2016; Gilbert & Whitlock, 2017; Hadfield, 2016; Schiffers et al., 2014) and several studies have framed this concept in terms of the spatial autocorrelation of the environment (Hadfield, 2016; Urban, 2011). Spatial autocorrelation describes the similarity of observations from nearby locations and can be quantified, for example, using Moran’s I (Moran, 1950). Consider the maps of environmental heterogeneity shown in Figure 1A. When the environment that gives rise to spatially varying selection across a species’ range exhibits high spatial autocorrelation (e.g. the right-hand map in Figure 1A), selection pressures may be similar over large areas and changes in environment over space will tend to be gradual. On the other hand, when the environment exhibits weak autocorrelation (e.g. the left-hand map in Figure 1A), regions experiencing idiosyncratic selection will be comparatively small and selection pressures may change rapidly over space. Of course, other factors such as variation in population density, the magnitude of environmental variation and the scale of dispersal will also affect the outcomes of spatially varying selection. All else being equal, though, a species with restricted migration will tend to evolve the strongest local adaptation if agents of selection exhibit high levels of spatial autocorrelation.

**Figure 1.**
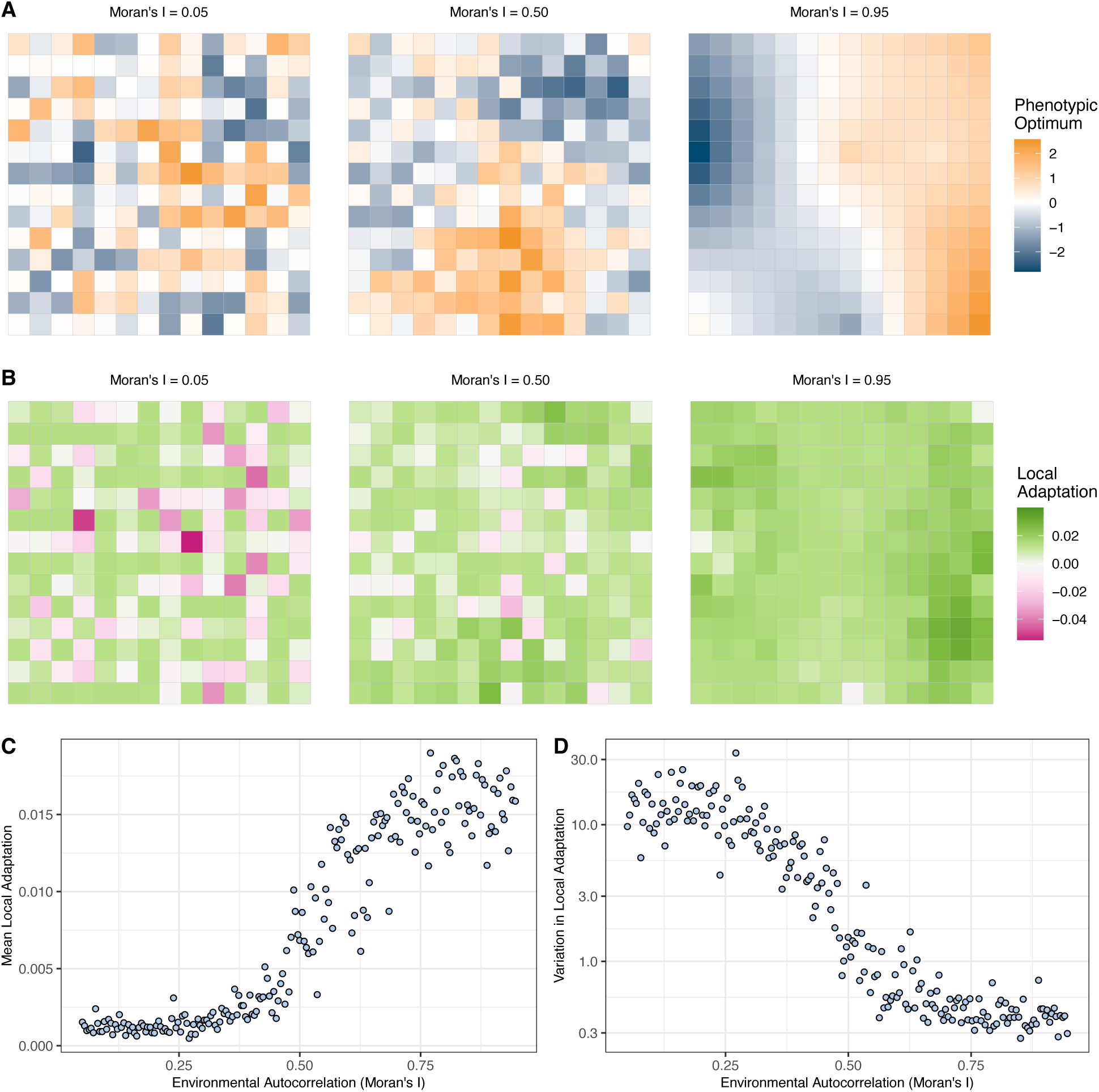
The pattern of environmental heterogeneity influences the outcomes of spatially varying selection. A) Three examples of environmental heterogeneity with similar distributions of phenotypic optima ranging from low autocorrelation on the left to high autocorrelation on the right. B) The pattern of local adaptation that evolved on the landscapes shown in panel A. C) The average local adaptation that arises as a function of Moran’s I across 200 maps of environmental variation. C) The coefficient of variation for local adaptation across the 200 maps. The simulation results shown are for cases with mean *F_ST_* ~ 2% and moderate stabilising selection.

It is likely that the spatial pattern of selection across a species’ range affects the genetic architecture underlying local adaptation. In simple two-patch models, the relative balance of selection and migration influences the number of alleles underlying local adaptation, their effect sizes and rates of allelic turnover (Yeaman & Whitlock, 2011). In continuous, linearly varying landscapes, frequencies of alleles contributing to locally adaptive traits are expected to vary in relation to their effect sizes and the proximity of local trait means to phenotypic optima (Polechová & Barton, 2015). However, such models may not fully predict the patterns of genetic variation expected in complex landscapes where the relative balance of selection and migration vary across space. Previous studies have examined the effects of landscape structure on the genetics of local adaptation in single locus models (Forester et al., 2016) or in models of population expansion (Gilbert & Whitlock, 2017; Schiffers et al., 2014), but it is unclear how the polygenic architecture of local adaptation will be influenced by the spatial pattern of environmental variation.

In this paper, I examine patterns and the genetic bases of local adaptation in complex landscapes. Following previous studies, I cast spatial patterns of environmental variation across a species’ range in terms of spatial autocorrelation. With population genetic simulations, I examine the patterns and genetic architectures of local adaptation that evolve in environments that vary in their degree of spatial autocorrelation. These simulations show ways that the pattern of environmental variation across a species’ range can influence the genetic variation underlying local adaptation. Finally, I analyse empirical data from a large-scale experiment in lodgepole pine and find evidence suggesting a link between spatial autocorrelation in climatic/environmental variation with the strength of local adaptation in a natural system. Taken together, the results of this study highlight the importance of considering the spatial pattern of environmental variation in studies of local adaptation.

## Results and Discussion

### Simulating local adaptation to spatially heterogeneous environments

To further understand the effects that the spatial pattern of environmental variation can have on patterns of local adaptation, I constructed a simulation model of spatially varying selection. I used forward-in-time population genetic simulations in SLiM (4.1; Haller & Messer, 2023) modelling a 2-dimensional stepping-stone metapopulation of 196 demes (i.e. a 14×14 grid). Migration was restricted to adjacent demes with rates of gene flow that resulted in pronounced population structure with clear isolation-by-distance (Figure S1). Spatially varying selection was modelled as stabilising selection, where each deme (*d*) had a particular phenotypic optimum (*θ_d_*), i.e. an individual in deme *d* had higher fitness if its phenotype was close to *θ_d_*. The strength of stabilising selection was set such that an individual with the optimal phenotype for the deme with the most negative optimum translocated into the deme with the most positive optimum would experience a 50% reduction in fitness (strong selection) or a 25% reduction in fitness (moderate selection). I used a quantitative trait model to study local adaptation, because it is thought that the traits involved in local adaptation are generally polygenic (Savolainen et al., 2013). Additionally, there is evidence that the genetic basis of local adaptation can involve alleles that have spatially antagonistic fitness effects as well as conditionally neutral effects (Anderson et al., 2013) and both kinds of effects can arise in a quantitative trait model given variation in genetic backgrounds and environments. I constructed a set of 200 maps of normally distributed environmental variation that varied in the degree of spatial autocorrelation (three examples are shown in Figure 1A). I quantified spatial autocorrelation in the environment using Moran’s I, which varied from 0.05 (weak autocorrelation) to 0.95 (strong autocorrelation) in the maps I constructed. The maps of environmental variation were used to specify phenotypic optima in individual simulations.

The effects of environmental structure on patterns of local adaptation that evolved in simulations were profound. I measured local adaptation in simulated populations by comparing an individual’s fitness at “home” versus “away” following the method outlined by (Blanquart et al., 2013). Using this method, the observed local adaptation for each deme (*LA*) in the metapopulation was computed (e.g. Figure 1B). As expected, the mean local adaptation across populations (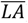) increased with the degree of spatial autocorrelation in the environment (Figure 1C), consistent with Hadfield (2016). This result held over various levels of gene flow and strengths of stabilising selection (Figure S2). Varying the rate of gene flow and/or the strength of selection had an effect on the level of local adaptation that arose in a given case, but a pattern of increasing 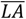 with Moran’s I was always observed (Figure S2). In natural populations, it is likely that multiple aspects of environmental variation will induce spatially varying selection. In cases where simulated populations had multiple traits subject to selection due to different aspects of environmental variation, the trait corresponding to the more autocorrelated environment exhibited greater local adaptation (Figure S3).

The spatial pattern of environmental variation did not just affect the average level of local adaptation, though, it also had a large influence on the variation in local adaptation across the landscape (Figure 1B). The coefficient of variation in local adaptation [*CV*(*LA*)] across the landscape decreased rapidly with increasing autocorrelation (Figure 1D, S2B). When environmental variation was weakly autocorrelated, the *CV*(*LA*) was as much as 30x higher than for more highly autocorrelated environments (Figure 1D). Variation in the degree of local adaptation across a species range is understudied in the population genetics literature but has important implications (see below).

At a finer scale, environmental variation in the immediate vicinity of a particular deme predicted its level of local adaptation and genetic variation, as predicted by theory (Barton, 1999; Guillaume & Whitlock, 2007; Slatkin, 1978). Demes that were surrounded by populations with highly similar phenotypic optima evolved greater local adaptation than demes bordering more dissimilar environments (Figure S4A). This was particularly evident when the overall landscape was weakly autocorrelated (Figure S4A), presumably because in highly autocorrelated landscapes most demes are surrounded by similar environments. Furthermore, additive genetic variance (*V_A_*) for the trait under selection was highest in demes surrounded by dissimilar environments (Figure S4B), suggesting that gene flow among locally divergent populations has an effect of increasing genetic variability. Such a positive correlation between *V_A_* and local environmental heterogeneity has been reported in lodgepole pine (Yeaman & Jarvis, 2006). The magnitude of a species’ response to selection on a trait is expected to be proportional to *V_A_* (Falconer & MacKay, 1995), thus the pattern of environmental variation that local adaptation evolves under may influence how a species responds to changing environments.

### Environmental structure and the genetic architecture of local adaptation

The results so far demonstrate that the structure of the environment can have a clear impact on the patterns of local adaptation that evolve, but does it influence the genetic basis of that adaptation? Under a model of spatially varying stabilising selection, each polymorphism that affects the phenotypes under selection will influence local adaptation, but the extent of this will depend on its effect size, where it is present and its allele frequencies. For each polymorphism in a simulation, I quantified the contribution it makes to mean local adaptation (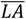) as follows. I shuffled the presence/absence of a particular polymorphism across the landscape, effectively erasing its contribution to local adaptation. Average local adaptation was then recalculated without the contribution of the focal polymorphism (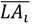). The relative contribution of the focal polymorphism to mean local adaptation across the landscape was then calculated as 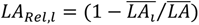. For example, a polymorphism with *LA_Rel_ ≈* 1.0 would be the basis of all local adaptation, while one with *LA_Rel_ ≈* 0.0 would have no effect. Note, *LA_Rel_* is not strictly a proportion (see Methods for details).

The distribution of locally adaptive effects varied in relation to the pattern of environmental variation (Figure 2A, S5). In environments exhibiting weak autocorrelation, polymorphisms that individually made a large contribution to local adaptation across the species’ range (*LA_Rel_* > 0.10) were largely absent and polymorphisms that made intermediate (0.01 < *LA_Rel_* < 0.10) and small contributions (*LA_Rel_* < 0.01) explained most of the local adaptation that evolved (Figure 2A). In environments that were more highly autocorrelated, polymorphisms with *LA_Rel_* > 0.10 made a substantial contribution to local adaptation alongside those with intermediate and small effects, particularly under strong stabilising selection (Figure S5). These general patterns held over different levels of gene flow (Figure S5).

**Figure 2.**
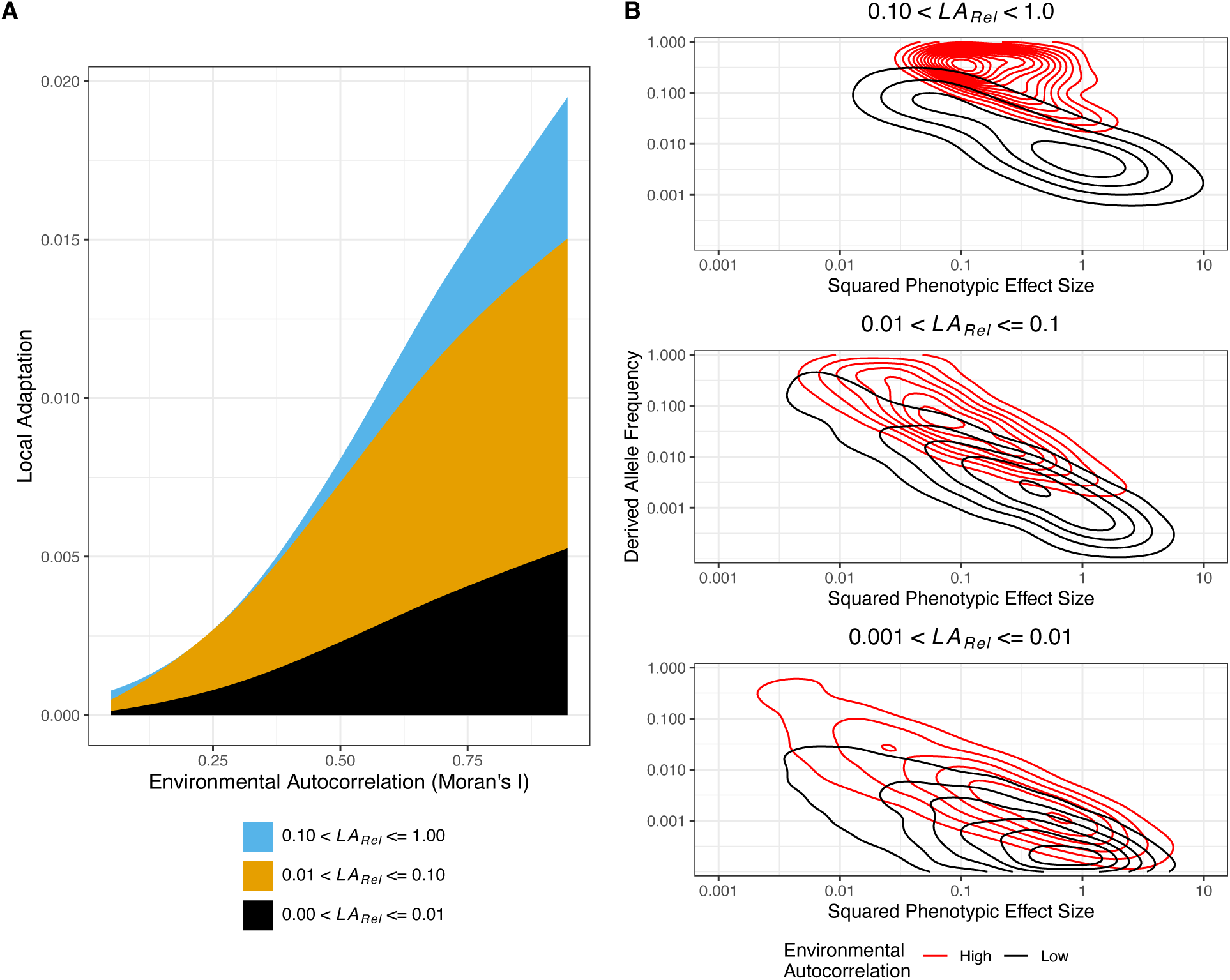
The genetic architecture of local adaptation is influenced by the structure of the environment. A) The proportion of total local adaptation explained by alleles that individually explain different amounts of local adaptation (*LA_Rel_*) varies as a function of environmental autocorrelation. Lines represent LOESS curves fit with a span parameter of 1.5. B) The mean allele frequencies compared to the squared effect sizes of polymorphisms that underly local adaptation differ depending on the pattern of the environment. The contour lines indicate regions with high densities of points. High autocorrelation refers to data from maps with the 50 highest values of Moran’s I. Low autocorrelation refers to data from maps with the 50 lowest values of Moran’s I. Results in both panels come from simulations with *F_ST_* = 0.02 and moderate stabilising selection.

Patterns of genetic variation underlying locally adaptive polymorphisms varied depending on the degree of spatial autocorrelation in the environment. In general, locally adaptive polymorphisms with similar *LA_Rel_* tended to have smaller phenotypic effects but larger allele frequencies in highly versus weakly autocorrelated environments (Figure 2B). In highly autocorrelated environments, alleles may readily spread among populations facing similar environmental challenges. However, when alleles with large phenotypic effects spread among neighbouring demes, they may cause individuals to overshoot their respective phenotypic optima, so the alleles that are maintained may tend to have smaller phenotypic effects. In weakly autocorrelated environments, on the other hand, genes flowing from one location to another have a much greater chance of encountering highly divergent environments, preventing locally adaptive alleles from spreading across wide regions. In such cases, phenotypic effects may need to be large for locally adaptive mutations to withstand the swamping effects of gene flow. These general patterns were observed with both strong and moderate selection (Figure 2B, S6A) as well as over varying levels of gene flow (Figure S6A). Indeed, the patterns of allele frequency versus phenotypic effect still held when looking at absolute effects on local adaptation, though they were much less pronounced (Figure S6B).

Characterising the genetic basis of local adaptation is important, and researchers generally attempt to do so using one of two strategies; by comparing phenotypic variation for traits important for local adaptation to genetic variation (i.e. genome-wide association studies, GWAS) or environmental variation to genetic variation (i.e. genotype-environment association analysis, GEA analysis)(reviewed in Lasky et al. 2023). Since patterns of genetic variation underlying local adaptation can differ depending on the pattern of environmental variation, statistical power to identify the genetic basis of local adaptation will likely vary for different aspects of the environment. To demonstrate this, I performed GWAS on phenotypes for 1,000 randomly chosen individuals from the simulations and corrected for population structure using the kinship matrix. Figure S7 shows that the −log_10_(*p*-values) for alleles that contribute similar levels of local adaptation tend to be smaller (i.e. there is less power) for high versus low autocorrelation environments. Some of this difference may partially be due to the population structure correction procedure (see below), but it demonstrates that the pattern of environmental variation that gave rise to local adaptation can affect the ability to study the genetics of that adaptation.

There are numerous factors that may interact with the pattern of spatially varying selection to shape the genetics of local adaptation that I did not explore here. The degree of genetic redundancy in relevant traits, distribution of phenotypic effect sizes, mutation rates and patterns of dispersal can all influence the genetics of local adaptation (e.g. Láruson et al., 2020; Yeaman, 2013; Yeaman & Whitlock, 2011). Follow up studies looking at how such factors influence the genetic basis of local adaptation in differently structured environments are needed. However, the results from the simulations should provide researchers seeking to characterise the genetic basis of local adaptation with useful intuition.

### The evolution of local adaptation: maladaptation and allelic turnover

In heterogeneous environments certain polymorphisms may have a net effect of reducing local adaptation across a species’ range. All populations will harbour such locally maladaptive alleles, because any new mutation that increases distance between an individual’s phenotype and the local optimum will reduce local adaptation even if only by a small amount. By summing the effects of all polymorphisms with *LA_Rel_* < 0 across a simulation, I obtained a measure of the cumulative local maladaptation across a meta-population. Note that the cumulative local maladaptation is analogous to “migration load”. All simulations exhibited some degree of maladaptation regardless of the level of autocorrelation, but the cumulative effects of maladaptive alleles were always higher under weakly versus highly autocorrelated environments (Figure 3A, S8B). Increasing the rate of gene flow increased the degree of maladaptation and increasing the strength of selection decreased it (Figure S8B).

**Figure 3.**
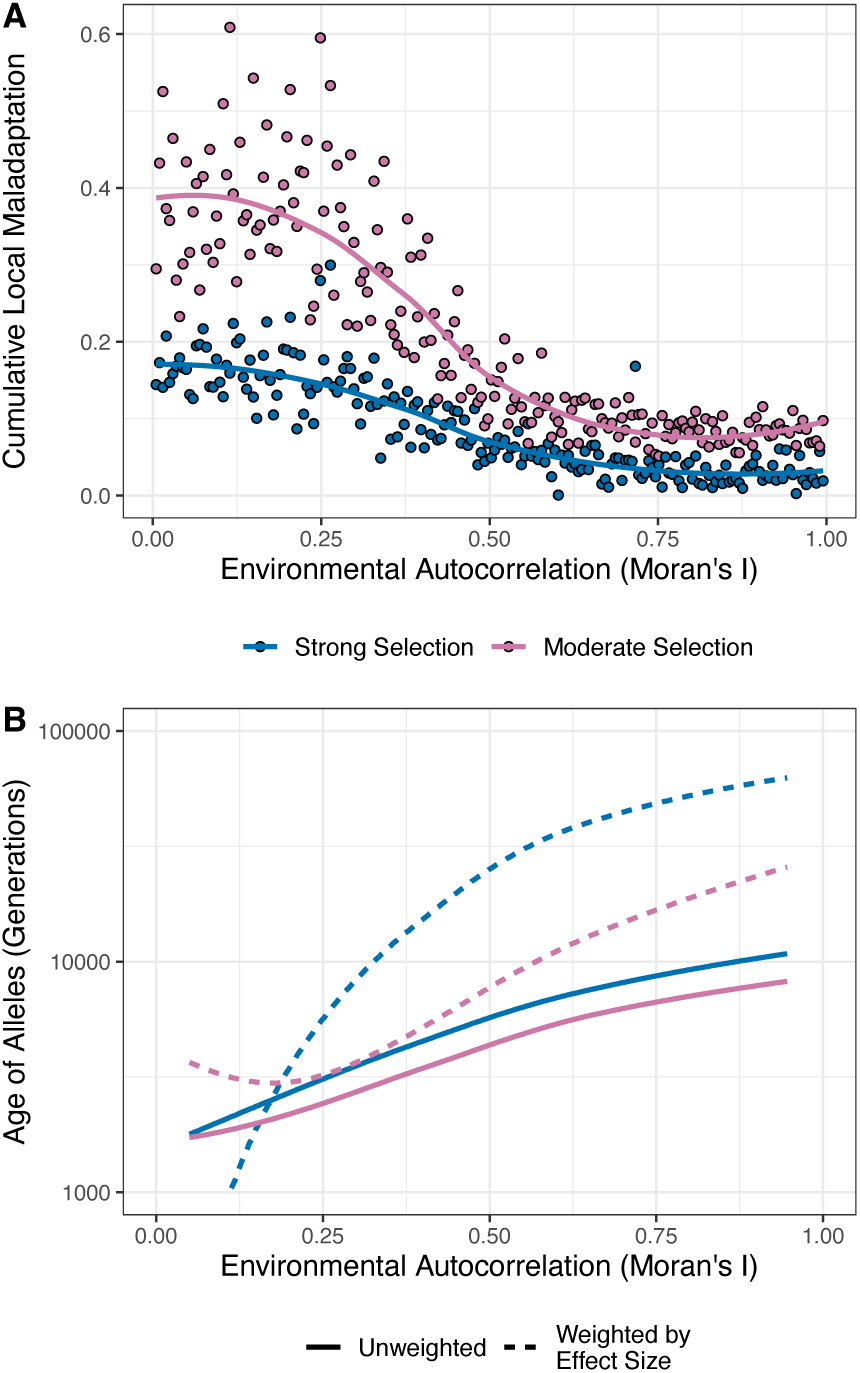
Species-wide maladaptation and the age of locally adaptive alleles are influenced by the pattern of environmental variation. A) Cumulative local maladaptation, the summed effects of all polymorphism that have a net negative effect on local adaptation across a simulated species’ range, decreases with increasing autocorrelation. Points represent individual simulations and lines represent LOESS curves fit with a span parameter of 1.5. B) Alleles underlying local adaptation tend to be older when the environment is more highly autocorrelated. Lines represent LOESS curves fit to the data treating all polymorphisms equally, or by giving higher weight to polymorphisms with greater effect size.

Regardless of the pattern of environmental variation in a simulation, levels of local adaptation had been maintained at a steady state for many generations before they were sampled (Figure S9). However, the average age of alleles underlying local adaptation increased with increasing autocorrelation in the environment (Figure 3B). Furthermore, weighing the average allele age within a simulation by effect size, the increase in allele age with spatial autocorrelation was even more pronounced (Figure 3B). Thus large effect alleles, in particular, are maintained for longer times in more highly autocorrelated landscapes. Taken together, these results demonstrate that the rate of allelic turnover is greater for more weakly autocorrelated environments.

### Local adaptation, environmental autocorrelation, provenance trials and lodgepole pine

The simulation results clearly show how the pattern of environmental variation across a species range may influence the evolution of local adaptation and the genetics underlying it. In natural populations, if the relative strength of selection acting on different aspects of environmental variation is unknown, we should perhaps predict that local adaptation will be strongest when the environment is highly autocorrelated. Despite such strong predictions, though, empirical evidence that patterns of local adaptation coincide with spatially autocorrelated features of the environment is lacking (Siepielski et al., 2013). Spatial autocorrelation in patterns of biotic interactions can explain a large proportion of variation in trait differentiation among populations in several species (Urban, 2011), but such variation is not necessarily locally adapted. Local adaptation has been demonstrated in many forest tree species using provenance trials (Leites & Benito Garzón, 2023), but not to test the prediction that the pattern of environmental variation influences local adaptation.

Provenance trials involve planting multiple populations of a species in numerous common gardens to assess how “transfer distance”, the distance between home and the common garden, affects productivity (reviewed in Wadgymar et al., 2022). However, the structure of provenance trials is such that the methods I used to quantify local adaptation in the simulations above are not necessarily applicable. To demonstrate how provenance trial data could be used to quantify local adaptation, I conducted *in silico* provenance trials on the simulations (e.g. Figure 4A). As expected, the slope of fitness on transfer distance in provenance trials is increasingly negative with increasing autocorrelation (Figure 4B, S10) and strongly negatively correlated with mean local adaptation (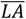) in simulations (Figure 4C, S10). Provenance trials may, thus, contain information that is useful for assessing whether the pattern of environmental across a landscape is important in shaping local adaptation.

**Figure 4.**
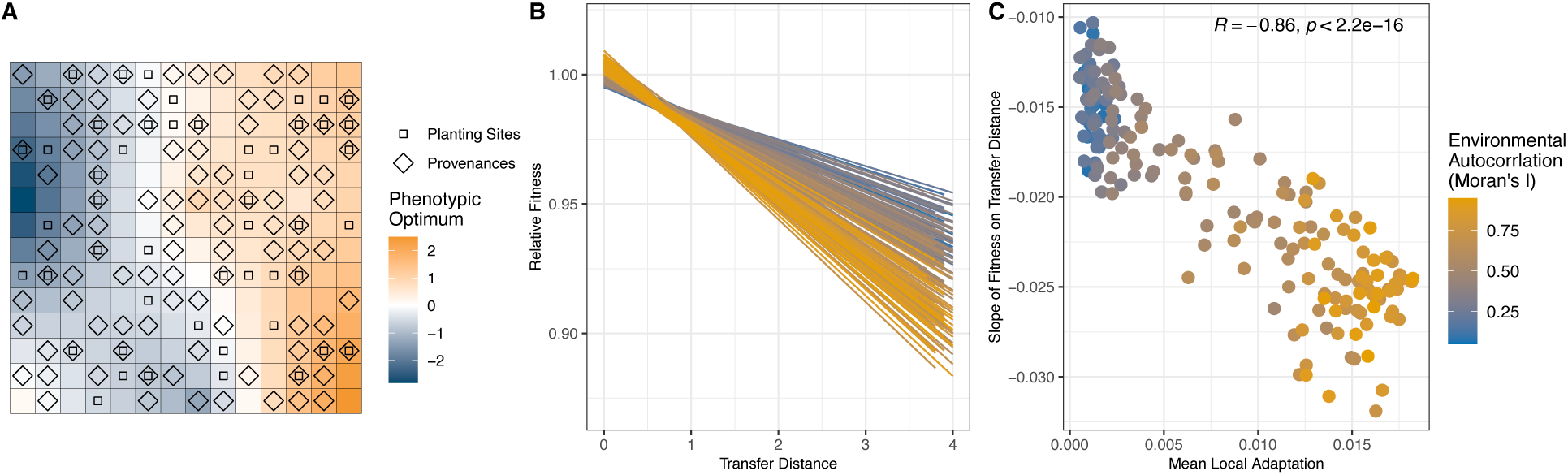
Comparing the results from a simulated provenance trial to measures of local adaptation. A) A map of a provenance trial conducted on a simulated population showing the locations of planting sites and provenances. B) Linear regressions of relative fitness on environmental transfer distance for landscapes with differing levels of environmental autocorrelation. C) The slope of relative fitness on transfer distance compared to the mean local adaptation (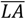) across simulated meta-populations. Spearman’s ρ and its *p*-value are shown inset in the panel C.

The Illingworth trial is an exceptionally large provenance trial established by the Ministry of Forestry in British Columbia, Canada in the 1970s to establish seed-transfer guidelines for the lodgepole pine (*Pinus contorta*) (Illingworth, 1978). Seeds were collected from 140 provenances from Northwestern North America and seedlings were planted in a set of 62 sites distributed across British Columbia (Figure 5A). Phenotypic data has been recorded for around 60,000 individual trees since the Illingworth trial began and previous studies have used this data to demonstrate clear local adaptation in lodgepole pine (Mahony et al., 2020; Wang et al., 2006). Given the geographic breadth of the Illingworth trial (Figure 5A), it represents a suitable dataset to test the prediction that spatial patterns of environmental variation influence the evolution of local adaptation.

**Figure 5.**
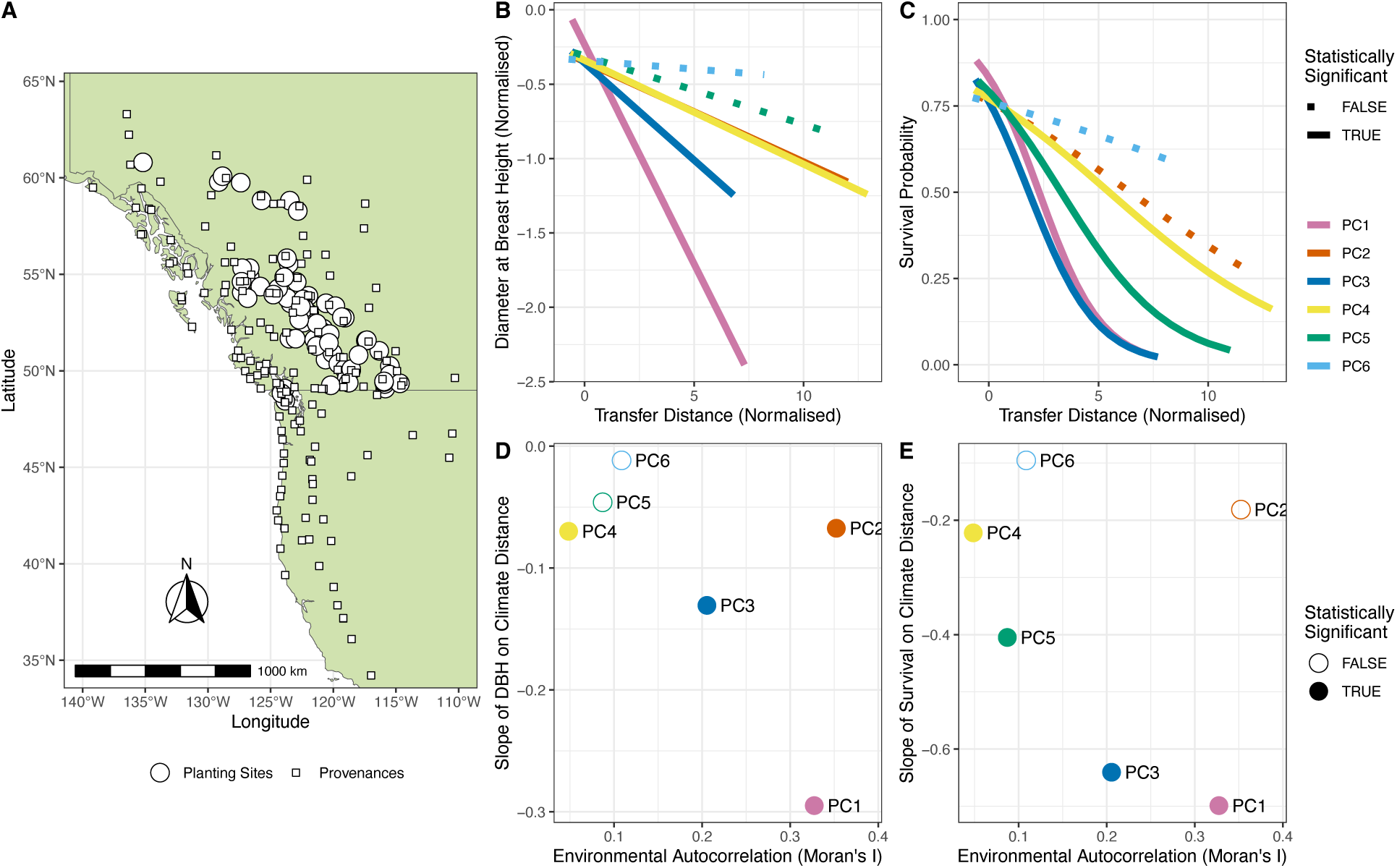
Analysis of local adaptation in lodgepole pine from the Illingworth provenance trial. Panel A) shows the map of provenances and planting sites in the Illingworth trials across the Northwest of North America. B) Fitted relationships between tree diameter at breast height (DBH) and transfer distance for 8 principal components. C) Survival probability as a function of transfer function for 8 environmental principal components. D) Linear regression coefficients for the relationships shown in B compared to degree of autocorrelation in the environment. E) Logistic regression coefficients for the relationships shown in C compared to the degree of spatial autocorrelation in the environment. Statistical significance was assessed at ɑ = 0.05 after correcting for multiple comparisons using the Dunn-Šidak method.

I analyzed data from the Illingworth trial using a mixed-modelling approach. Different aspects of climatic variation across the sites in the Illingworth trial are highly intercorrelated (Figure S11A), so I used principal components analysis to identify independent axes of climatic/environmental variation in the dataset. I restricted the analysis to the first 6 principal components (PCs), as each one explained at least 1% of the variation in the data and combined they explained 95% of the variation (Figure S11B). I then regressed tree diameter at breast height and survival measured after 20 years on transfer distance between planting site and provenances in PC-space (Figure 5B-C) (see Methods for details). For both diameter at breast height and survival, there were significant negative relationships predicting phenotypic variation from climatic PCs (Figure 5B-C). If diameter at breast height and/or survival are considered proxied for fitness, then the results shown in Figures 5B-C indicates local adaptation along several dimensions of climatic variability.

Patterns in the Illingworth trial data suggest that local adaptation in lodgepole pine is strongest when climatic/environmental variation is highly spatially autocorrelated (Figure 5C-D). For both phenotypes, PC1 was the strongest climatic predictor (Figure 5B-C). PC1 captures climatic/environmental differences between coastal and inland locations (Figure S12). Spatial autocorrelation for PC1 was also among the highest in the dataset (Figure 5D-E). For survival, there was a negative relationship between the regression slope and Moran’s I (Figure 5E), which is qualitatively similar to the analysis of simulated provenance trials (Figure 4B). However, I did not conduct a formal statistical test of the relationship between spatial autocorrelation and local adaptation because only 4 PCs exhibited statistically significant evidence for local adaptation so such an analysis would be underpowered. While this means the results are merely suggestive rather than concrete, they are in line with the prediction that spatial autocorrelation in the abiotic environment predicts the strength of local adaptation in natural populations.

### Population structure and the genetic basis of local adaptation

Knowing the genetic basis of local adaptation in natural systems would give us a better understanding of evolution, but may also be informative for conservation and management (Grummer et al., 2022). Many studies have used methods that associate environmental/phenotypic variation with genotypes or allele frequencies to characterise the genetic basis of local adaptation (Lasky et al., 2023). A large proportion of species exhibit a specific pattern of population structure termed “isolation-by-distance” (IBD) where genetic distance is positively correlated with geographic distance or measures of resistance based on features of the landscape (Jenkins et al., 2010). IBD can arise with restricted migration, when rates of gene flow are highest among parts of a species’ range that are in close proximity, though it can also reflect demographic histories such as past population expansion (Slatkin, 1993). Greater levels of local adaptation may evolve across a species’ range if the spatial pattern of environmental variation aligns with the opportunity for migration (Figure 1). Thus, stronger local adaptation is expected to arise when selection is co-linear or co-autocorrelated with patterns of population structure.

The relationship between population structure and the structure of the environment likely impacts our ability to study the genetic basis of local adaptation. It is well established that a pattern of IBD can confound the search for genes involved in local adaptation (Meirmans, 2012). Indeed characterising the genetic basis of local adaptation when the agents of selection are co-linear or co-autocorrelated with patterns of gene flow, termed “isolation by environment”, is particularly challenging (Wang & Bradburd, 2014). Many studies have analyzed genotype-environment associations (GEA) to characterize the genetic basis of local adaptation. Such association methods often treat population structure as a nuisance variable and various approaches are taken to correct for it. This is done for the statistical necessity of establishing a suitable null model (Meirmans, 2012). For example, latent factor mixed models (LFMMs) are widely used to conduct GEA analyses that correct for population structure (Caye et al., 2019; Frichot et al., 2013). However, Lotterhos (2023) recently found that the sensitivity of the LFMM method declined with increasing correlation between the environment and major axes of population structure. This all suggests that characterising the genetic basis of local adaptation is particularly difficult in the exact cases where local adaptation is expected to be strongest, when selection pressures and population structure are highly co-autocorrelated over space. Careful sampling strategies may alter the power of association methods (Lotterhos & Whitlock, 2015; Meirmans, 2015; Wang & Bradburd, 2014), but such strategies require *a priori* hypotheses about locally adapted traits, the agents of selection (e.g. Kreiner et al., 2022) and/or the genes involved (e.g. Fournier-Level et al., 2011).

### Implications for conservation management

A result from the simulations that was particularly striking is the heterogeneity in local adaptation that can arise under different patterns of environmental variation. The only difference between the simulations shown in Figure 1 is the pattern of environmental variation, yet the average level of local adaptation increased by a factor of roughly 5x and the coefficient of variation in local adaptation decreased by 30x when comparing the cases with the highest and lowest spatial autocorrelation in the environment. High heterogeneity in local adaptation across a species’ range may impact conservation interventions and population genetic analyses.

Any practical conservation intervention that uses average patterns of local adaptation to project performance under changing climates should carefully consider heterogeneity in local adaptation. In forestry, for example, seed-transfer guidelines based on average transfer functions from provenance trials may give a misleading picture of performance for some provenances if there is high heterogeneity in local adaptation. The applicability of a single transfer function would vary depending on how heterogeneous local adaptation is among the populations in question.

In recent years, population genetic analyses have been developed to identify parts of a species range that are particularly vulnerable to climate change (i.e. genomic offset; Rellstab et al., 2021). Such methods analyse present-day relationships between allele frequency and the environment to predict how species will fare given predicted patterns of environmental change. If the agents of local adaptation are highly spatially autocorrelated, neutral population structure may be partially aligned with gradients of selection, which could explain why the use of “adaptive” genetic markers and randomly chosen markers seem to perform equally well in some offset analyses (Fitzpatrick et al., 2021; Láruson et al., 2022; Lind et al., 2023). Violating the assumption of homogeneous local adaptation in offset analyses, for example, would likely introduce noise into predictions but could potentially lead to spurious results.

### Thinking about environmental structure when building hypotheses about local adaptation

Detailed prediction of the environmental variation that is relevant to patterns of local adaptation requires an understanding of a species’ life history and physiology. The fundamental factor underlying the evolution of local adaptation is the relative balance of selection and dispersal (see Introduction). However, the specific pattern of environmental variation across a landscape influences whether dispersing individuals are likely to encounter environments similar to those of their parents. Unlike dispersal or the strength of selection, which are hard to quantify, spatial autocorrelation in the environment is readily measurable. When seeking to characterise the genetic basis of local adaptation, studies comparing different aspects of environmental variation should consider the pattern of such variation when forming their hypotheses. For example, before comparing GEA results for the different *bioclimatic* variables from WorldClim, researchers could examine how these variables are distributed over space to form *a priori* hypotheses about factors underlying local adaptation. Of course, an aspect of the environment may be highly autocorrelated in space, but if the variation it exhibits does not correspond to varying selection pressures, then it is unlikely to be directly related to local adaptation. That local adaptation is predicted to be stronger with increasing autocorrelation in the environment does not imply that strong local adaptation cannot arise in highly heterogeneous environments or with little spatial autocorrelation. There are numerous examples of local adaptation to environmental heterogeneity that is not smoothly distributed in space. For example, heavy metal concentrations in mine tailings impose selection that is so strong it overwhelms the effects of gene flow (Jain & Bradshaw, 1966). It must be kept in mind that the patchy distributions of environmental variation across a landscape, as may be the case for heavy-metal rich soils, may be more or less autocorrelated from the perspective of a given species depending on its dispersal behaviour. In my simulations, I matched the granularity of dispersal with that of the environment (Figure 1A). For real species, considering environmental variation at a scale relevant to how species disperse is critical and recent population genetic advances may make estimating dispersal in natural populations much less time-intensive than previously (Bradburd & Ralph, 2019; Smith et al., 2023). Comparing patterns of species dispersal with patterns of variation in environmental variation that is plausibly relevant to selection may help identify the drivers of local adaptation in natural populations.

### Closing remarks

While it has been a long-standing expectation that the pattern of environmental variation (and particularly spatial autocorrelation) will influence the evolution of local adaptation (e.g. Hadfield, 2016; Levins, 1966), the simulation results and analysis of the lodgepole pine data should serve to emphasise how important the spatial pattern of climatic/environmental variation can be. The spatial pattern of environmental variation that a natural population has experienced will have likely shaped the evolution, current patterns and genetic underpinnings of local adaptation. Thus, it likely also influences how populations will respond to changes in the future.

## Materials and Methods

### Simulating spatially varying selection

To explore the effects of landscape structure on the outcomes of spatially varying selection, I constructed set of maps that exhibited varying degrees of spatial autocorrelation. Maps of normally distributed environmental heterogeneity were constructed using the midpoint displacement algorithm as implemented in the *NLMpy* package (Etherington et al., 2015). I simulated a 14×14 cell grid (i.e. landscape), specifying the desired level of autocorrelation to achieve a set of 200 maps, spanning the range of Moran’s I values from 0.05 to 0.95 (i.e. Moran’s I varied in increments of 0.0045). Simulated maps were rejected if the mean value across the landscape was less than 0.4 or greater than 0.6. This ensured that the the approximately normal distributions of environmental values across the landscape were roughly equivalent across maps.

Using SLiM v4.1 (Haller & Messer, 2023), I modelled a 2-dimensional stepping-stone meta-populations with 196 demes (i.e. a 14×14 grid). Each deme contained 100 diploid individuals for total meta-population size of 19,600. Migration occurred between adjacent demes in the four cardinal directions except for populations at the range edge where migrants only moved back into demes they were connected to. Migration rates were set at 0.07, 0.035 or 0.0175, leading to population-wide neutral *F_ST_* values of 0.02, 0.05 and 0.10, respectively (Figure S1A). Each diploid individual had a 10Mbp long genome that recombined at a constant rate of *r*=1 × 10^−7^. When modelling a single trait, mutational effects were distributed as *N*(0,1) and occurred at random along the sequence at a rate of *μ* = 10^−10^, corresponding to a mutational variance of 0.001 for the trait subject to stabilising selection. When modelling two traits, the mutation rate was the same, but effects were modelled as multivariate normal with means of 0, variances of 1 and covariances of 0 (i.e. mutational effects for the two traits were independent). A diploid individual’s phenotype for a given trait was the additive combination of the effects on that trait for the alleles the individual possessed (i.e. mutations were semidominant). Spatially varying stabilising selection was modelled using the maps of environmental heterogeneity to specify the distribution of phenotypic optima across the landscape. An individual’s relative fitness *W_i_* was calculated using the standard expression for Gaussian stabilising selection (Walsh & Lynch, 2018):

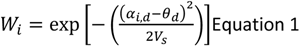

where V_S_ is the variance of the Gaussian fitness function, α_i_ is the phenotype of the i^th^ individual in deme d, and *θ*_d_ is the phenotypic optimum of deme d. When modelling stabilising selection in cases with two traits, an individual’s fitness was calculated as follows:

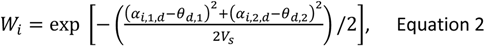

where α_i,1,d_ and α_i,2,d_ are the values for traits 1 and 2 for individual i in deme d, respectively, and *θ*_d,1_ and *θ*_d,2_ are the phenotypic optima for traits 1 and 2, respectively. In effect, an individual’s relative fitness in this 2-trait model is the average of the marginal finesses for each trait.

To achieve an equilibrium of migration, selection and drift, meta-populations evolved for 100,401 generations. Initially, meta-populations evolved under stabilising selection with an optimum of 0 in all demes. After 400 generations, the landscape was altered to one of the 200 maps of environmental heterogeneity and kept in that state for a further 100,000 generations. At the end of the simulation, phenotypes of each individual in each deme were recorded as well as the genealogical history of the meta-population stored as a tree-sequence. PySlim, tskit and msprime packages (Baumdicker et al., 2022; Haller et al., 2019) were used to work with the output tree-sequence files. To calculate Weir and Cockerham’s F_ST_, neutral mutations were added to the simulated population using PySlim at a rate of 10^−8^/bp.

### Analyzing simulated data

Local adaptation was quantified for each deme using the “home-versus-away” (HA) method outlined by Blanquart et al. (2013). Specifically, each individual’s fitness was quantified in its home deme and every other possible location on the landscape. The mean local adaptation was calculated in each deme as the mean difference in fitness between home and away conditions across all individuals. For each deme *d* not on the edge of the simulated landscape, I quantified local heterogeneity in the landscape as the mean sum of squares between the focal deme’s environment and that of the four adjacent demes (in the cardinal directions). For each deme, across the *n* polymorphisms that affected the phenotype additive genetic variance for the trait was calculated as 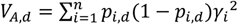, where *p_i_*_,*d*_ is the allele frequency of SNP *i* in deme *d* and *γ_i_* is the phenotypic effect of SNP *i*.

The contribution of individual SNPs to local adaptation was quantified as follows. For each polymorphism that affected the trait(s) under selection, the presence/absence of the allele in different haplotypes in different demes can be represented as a vector of 1s and 0s. By shuffling this vector, the contribution of this polymorphism to local adaptation is effectively erased, while keeping its contribution to additive genetic variance across the species’ range constant. For polymorphism *l,* I recomputed all phenotypes for all individuals after shuffling allele frequencies and re-quantified local adaptation as 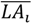. The relative contribution of the focal polymorphism to local adaptation is calculated as:

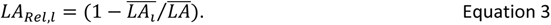

Note that *LA_Rel_* is not strictly a proportion, as epistasis for fitness that arises in models of stabilising selection means that the 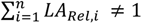 for the *n* SNPs that affect phenotypes. Furthermore, alleles that have a net negative effect on local adaptation (i.e. they are locally maladaptive) will have negative *LA_Rel_* values. Indeed, the total amount of local maladaptation in a meta-population was calculated as the additive combination of all polymorphisms with negative *LA_Rel_*.

Provenance trials were conducted on simulated data by sampling a set of 50 “planting sites” and a set of 100 “provenances”. The relative fitness of each provenance was computed in each of the 50 planting sites. The absolute difference in phenotypic optimum for each provenance and each planting site was used as environmental distance. Using *lme4* in R, I fitted a linear mixed model regressing relative fitness on environmental distance with provenance as a random effect, with slopes and intercepts varying across provenances.

I combined results across the 50 simulations with the lowest levels of spatial autocorrelation (weak autocorrelation) and the 50 simulations with the highest autocorrelation (high autocorrelation) and examined the relationship between allele frequency and phenotypic effect sizes for the alleles underlying local adaptation.

For each simulation, I randomly sampled 1,000 individuals from the landscape and I recording their trait values as well as neutral and phenotype affecting polymorphisms. I applied LD-pruning (with a threshold of *r^2^* < 0.2) to the neutral SNPs and used these data to infer the kinship matrix using PLINK (v2; Chang et al., 2015). I performed association studies on the phenotype affecting SNPs from individual simulations with GEMMA (v0.98.5; Zhou & Stephens, 2012), using the inferred kinship matrix as a random effect to account for population structure.

### Analysis of data from the Illingworth trial

ClimateNA (Wang et al., 2016) was used to extract climatic data for each location in the Illingworth Trial. Across the locations in the Illingworth trial, many aspects of climatic/environmental variation are highly inter-correlated (Figure S11B). Because such inter-correlation would make it difficult to tease apart the effects of individual aspects of climatic/environmental variation on local adaptation, I conducted a principal components analysis (PCA) to separate the variation onto independent axes. I restricted the analysis to the first 6 principal components as these explained 95% of climatic variation. Diameter at breast height (DBM) and tree height exhibit a strong positive correlation (Pearson’s *r* = 0.9), so analyses were restricted solely to DBM. Trees that were dead or dying after 20 years were given a survival score of 0, living trees were scored a 1.

Phenotype and survival data after 20 years for individual trees from the Illingworth trial were analyzed using mixed models. Mean normalised DBH was modelled as a normally distributed variable using the *lme4* package and survival using a generalised linear mixed model with a “logit” link function using the *glmer* package. The normalised Euclidean distance between each individual’s provenance and planting site (i.e. transfer distance) in PC-space was used as a predictor in the model. Provenance, planting site and planting block within sites were included as having random effects on the slope and intercept of the relationship between phenotype and transfer distance.

Moran’s I was calculated for each principal component of climatic variation across provenances (using the *ape* package) incorporating a pairwise Haversine distance matrix as weights in the calculation.

## Data accessibility

All the code used to perform, analyse, and plot the results of simulations is available at https://github.com/TBooker/LocalAdaptationArchitechture. R scripts to analyse and plot the results of the Illingworth trial data are available at https://github.com/TBooker/LocalAdaptationArchitechture, but the raw data files were used by permission of the BC Ministry of Forestry.

## Acknowledgements

Thanks to the Aitken and Whitlock labs, Anna Bazzicalupo, Katie Lotterhos, Ailene Macpherson, Greg O’Neil, Monty Slatkin, Wouter van der Wijl and Sam Yeaman for discussions and/or comments on the manuscript. Thanks to Hazel Booker for late night chat sessions. The data from the Illingworth trial was compliments of Nick Ukrainetz, the lodgepole pine breeder with the Forest Improvement and Research Management branch of the BC Ministry of Forests. TRB was supported by a Bioinformatics Postdoctoral Fellowship from the Biodiversity Research Centre at the University of British Columbia.

## Supplementary Material

**Supplementary Figure 1.**
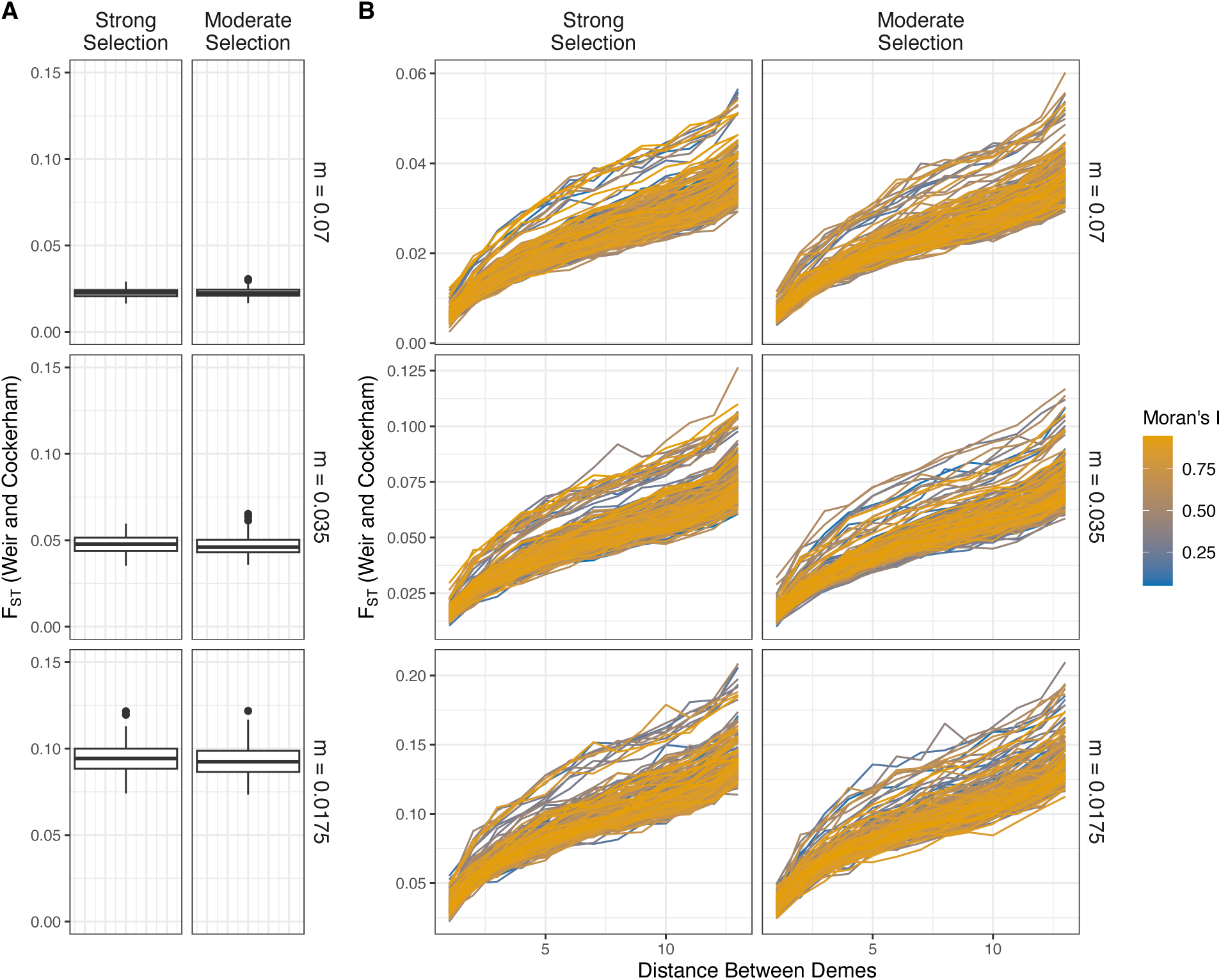
Overall *F_ST_* (panel A) and isolation by distance (panel B) in simulated populations. Note the varying y-axes in panel B. In the main text, *F_ST_* is used to refer to the panels of individual graphs. In panel A values from 200 independent simulations were used to construct the boxplot and in panel B individual simulations are shown as lines. Weir and Cockerham’s method for calculating *F_ST_*, as implemented in the *sci-kit-allel* Python package, was used.

**Supplementary Figure 2.**
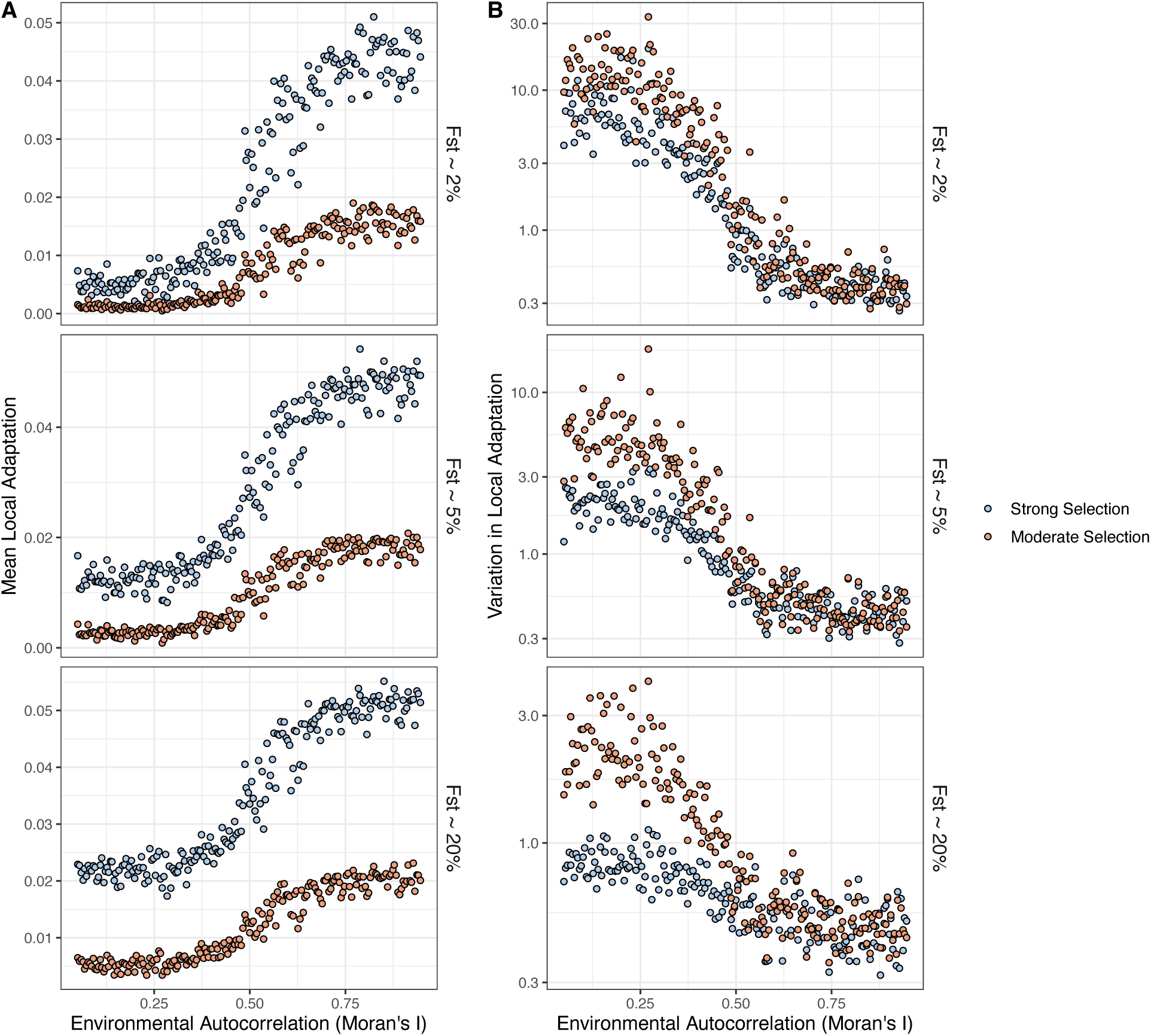
The average extent of local adaptation (panel A) and coefficient of variation in local adaptation (panel B) as a function of spatial autocorrelation in the environment from simulated datasets. The upper cell of each column is included in Figure 1 of the main text.

**Supplementary Figure 3.**
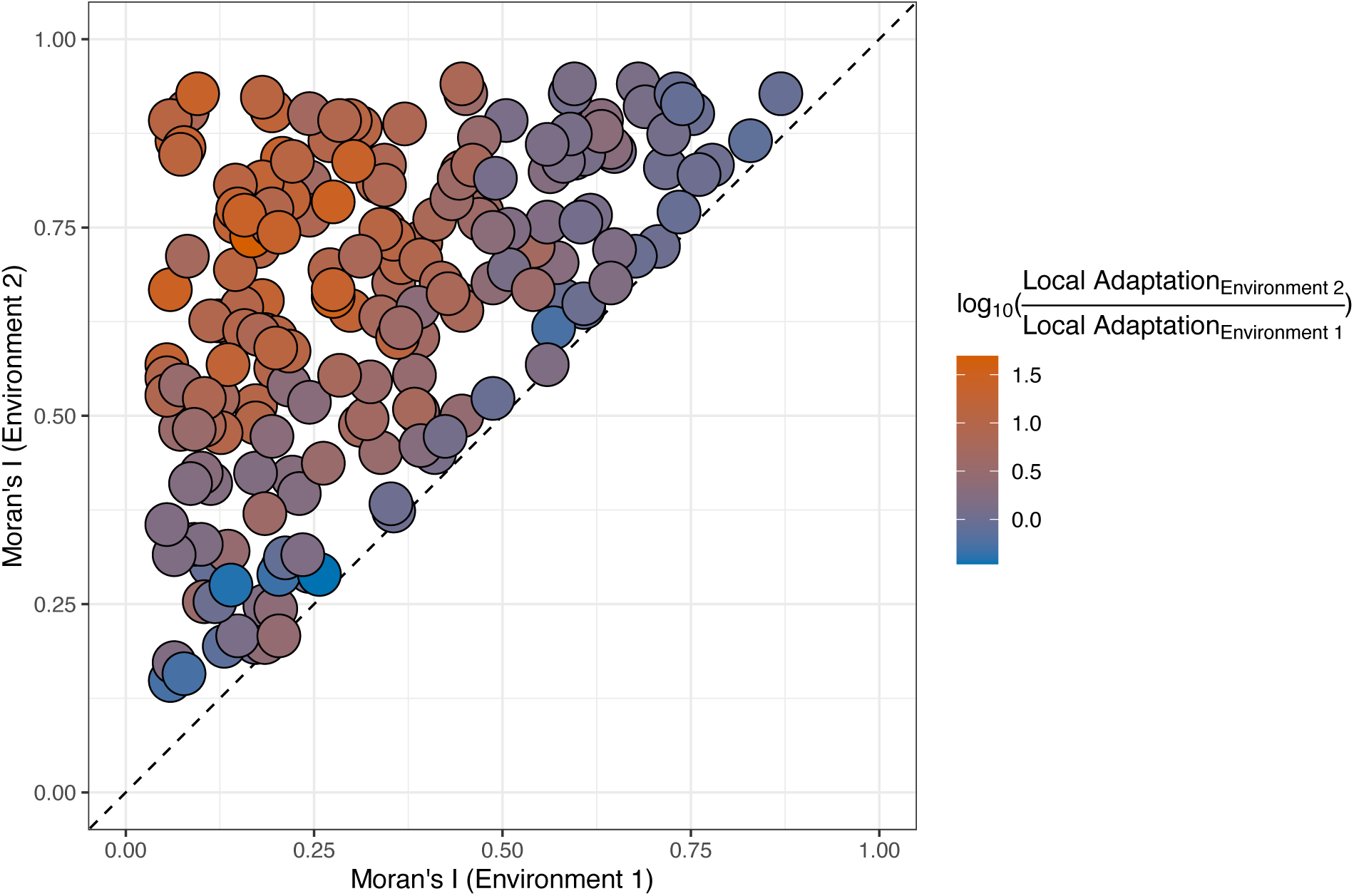
Comparison of local adaptation that evolves for two traits subject to spatially varying selection. Selection on each trait was dictated by distinct maps of environmental variation/phenotypic optima. The environment that exhibited the greater degree of spatial autocorrelation (as measured by Moran’s I) was designated “Environment 2”. The 1:1 line is shown for reference.

**Supplementary Figure 4.**
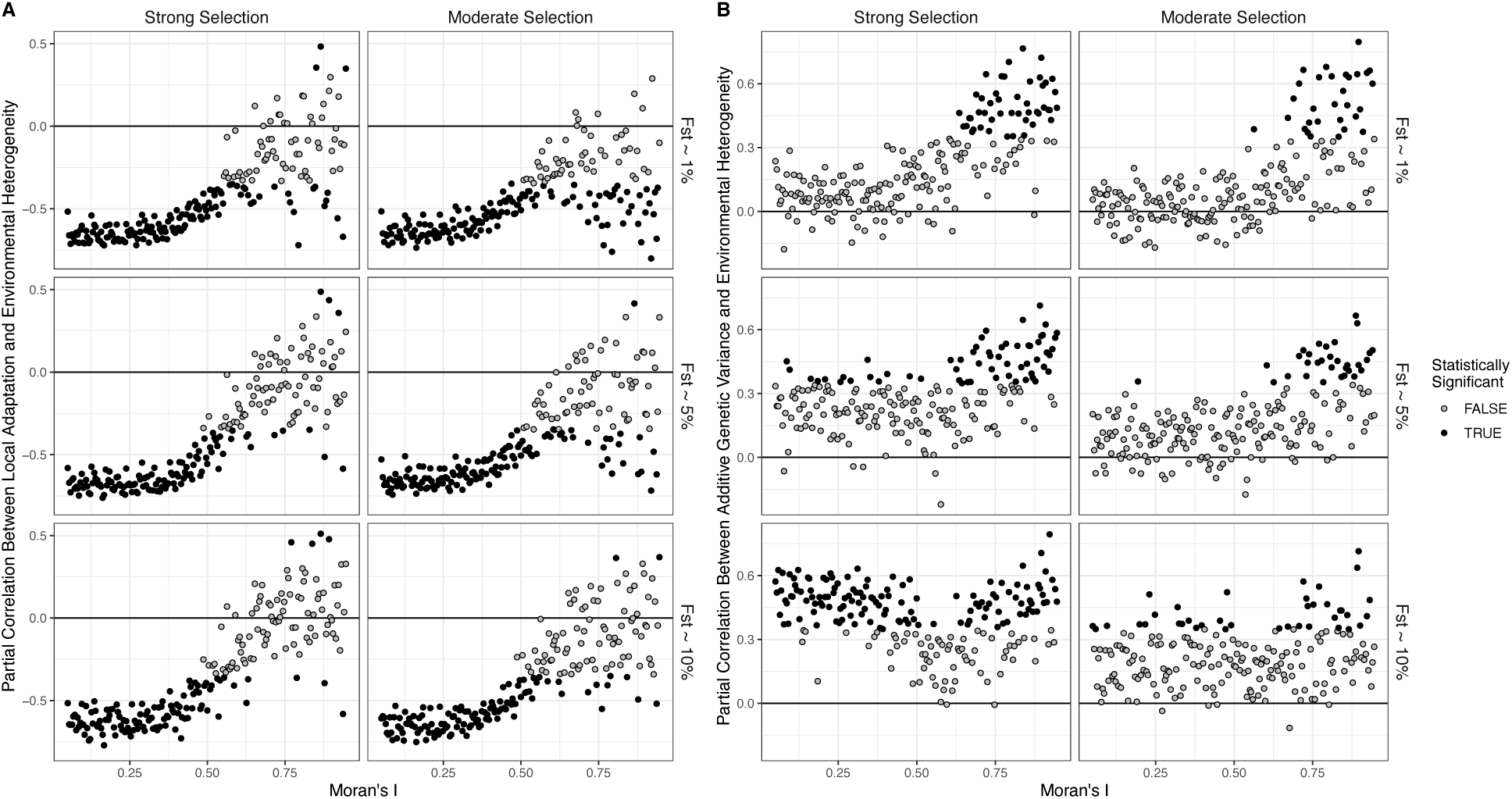
The effect of spatial autocorrelation in the environment on the correlations of local environmental heterogeneity with local adaptation and additive genetic variance. A) The partial correlation between local adaptation and local environmental heterogeneity, controlling for additive genetic variance. B) The partial correlation between additive genetic variance and environmental heterogeneity, controlling for local adaptation. Statistical significance was assessed after correcting for multiple comparisons. The solid black line indicates the statistical null expectation of 0.

**Supplementary Figure 5.**
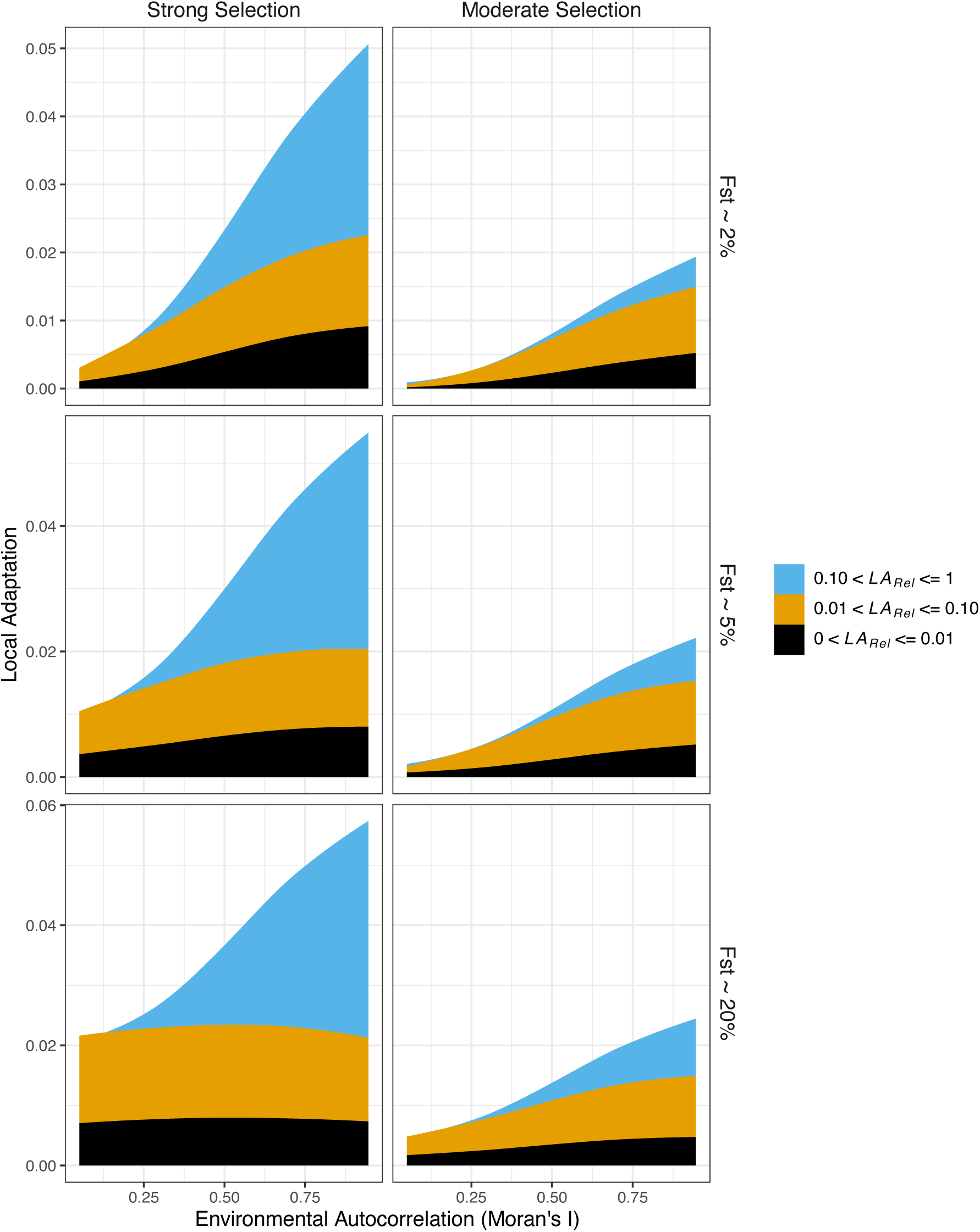
The distribution of locally adaptive effects as a function of spatial autocorrelation in the environment. The area shown was calculated across 200 independent simulations and smoothed using a LOESS regression with span 1.5. The upper right cell is included in Figure 2 of the main text.

**Supplementary Figure 6.**
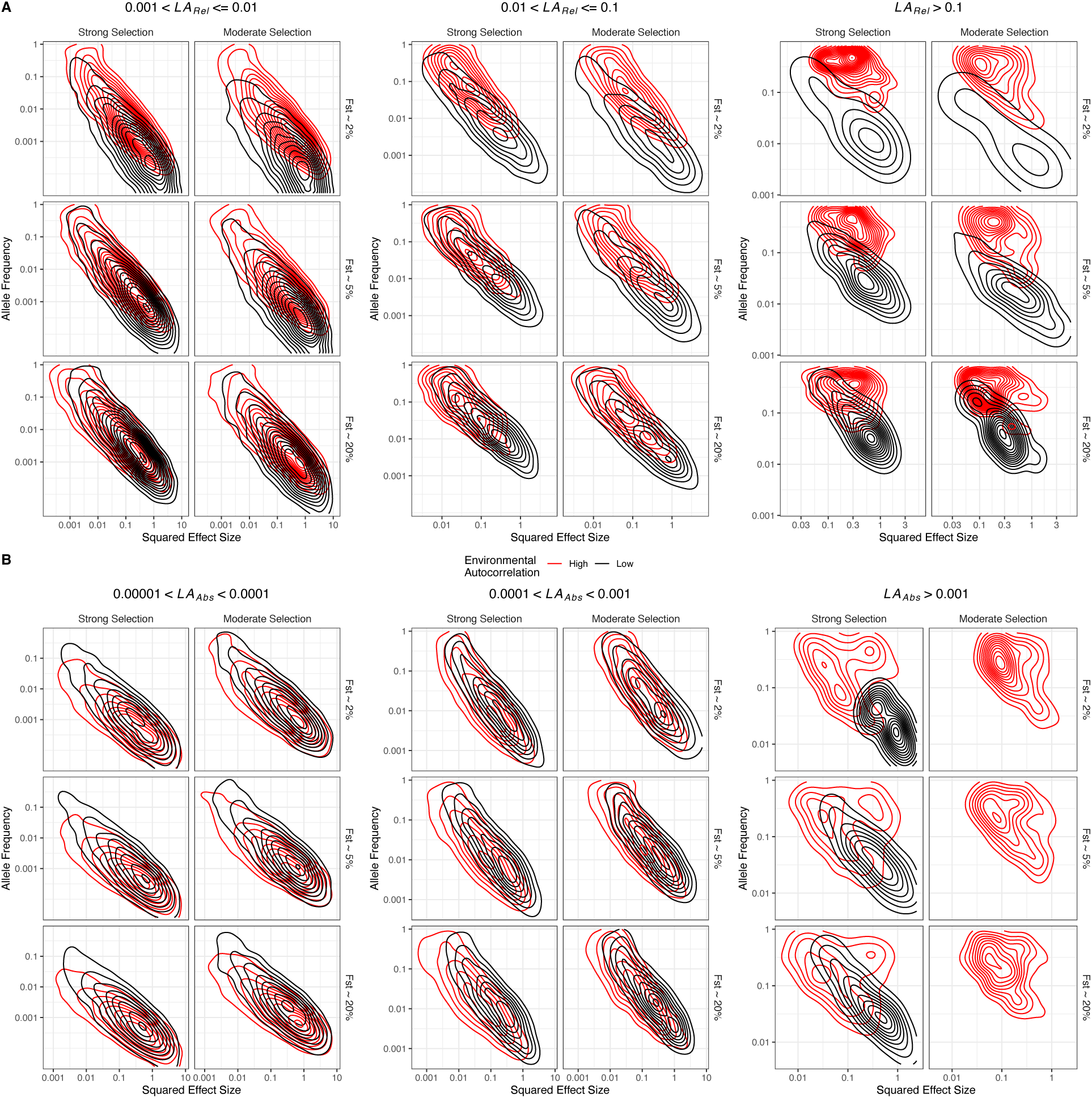
The relationship between allele frequency and the squared phenotypic effect size for polymorphisms that contribute varying degrees of local adaptation in either relative (panel A) or absolute terms (panel B).

**Supplementary Figure 7.**
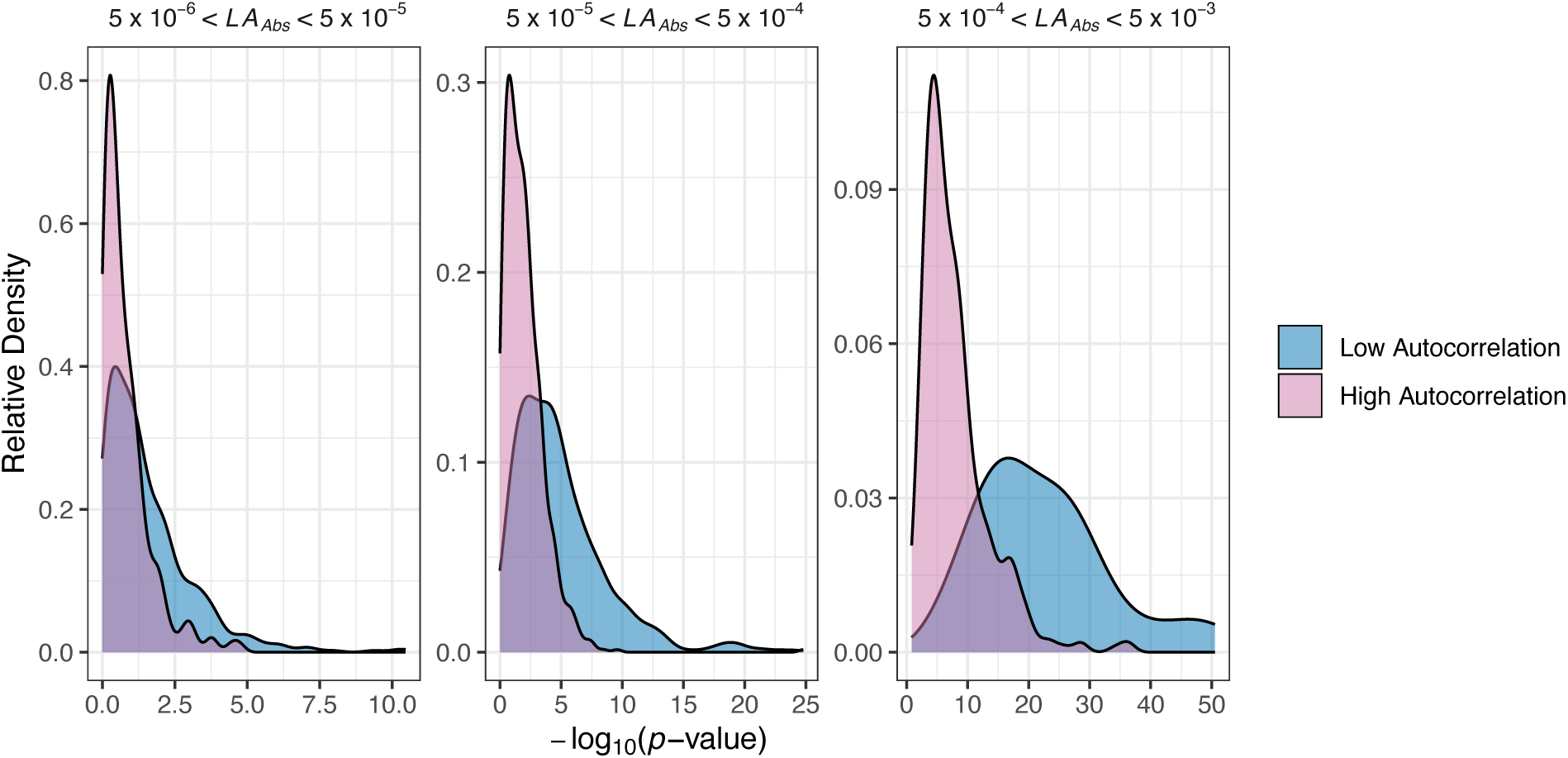
Results from a GWAS on 1,000 randomly chosen individuals from either high or low autocorrelation environments. Each panel compares the relative density of −log10(*p-*values) from a GWAS conducted on data from the 50 maps with the highest or lowest levels of spatial autocorrelation.

**Supplementary Figure 8.**
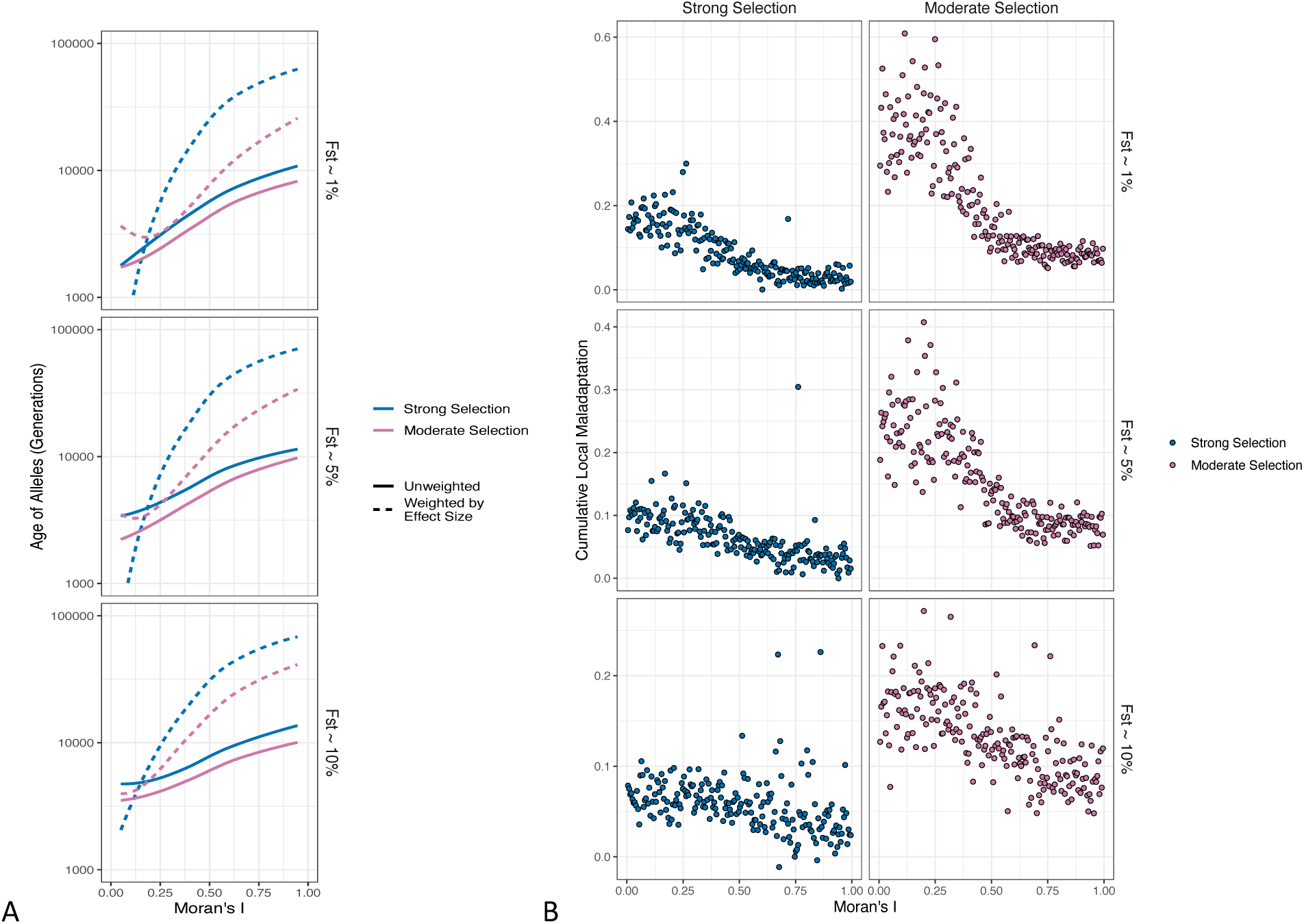
A) Cumulative local maladaptation as a function of spatial autocorrelation in the environment across all parameter combinations. B) The average age of locally adaptive alleles in meta-populations subject to spatially varying selection. The lines represent LOESS regression curves calculated with span parameters of 1.5.

**Supplementary Figure 9.**
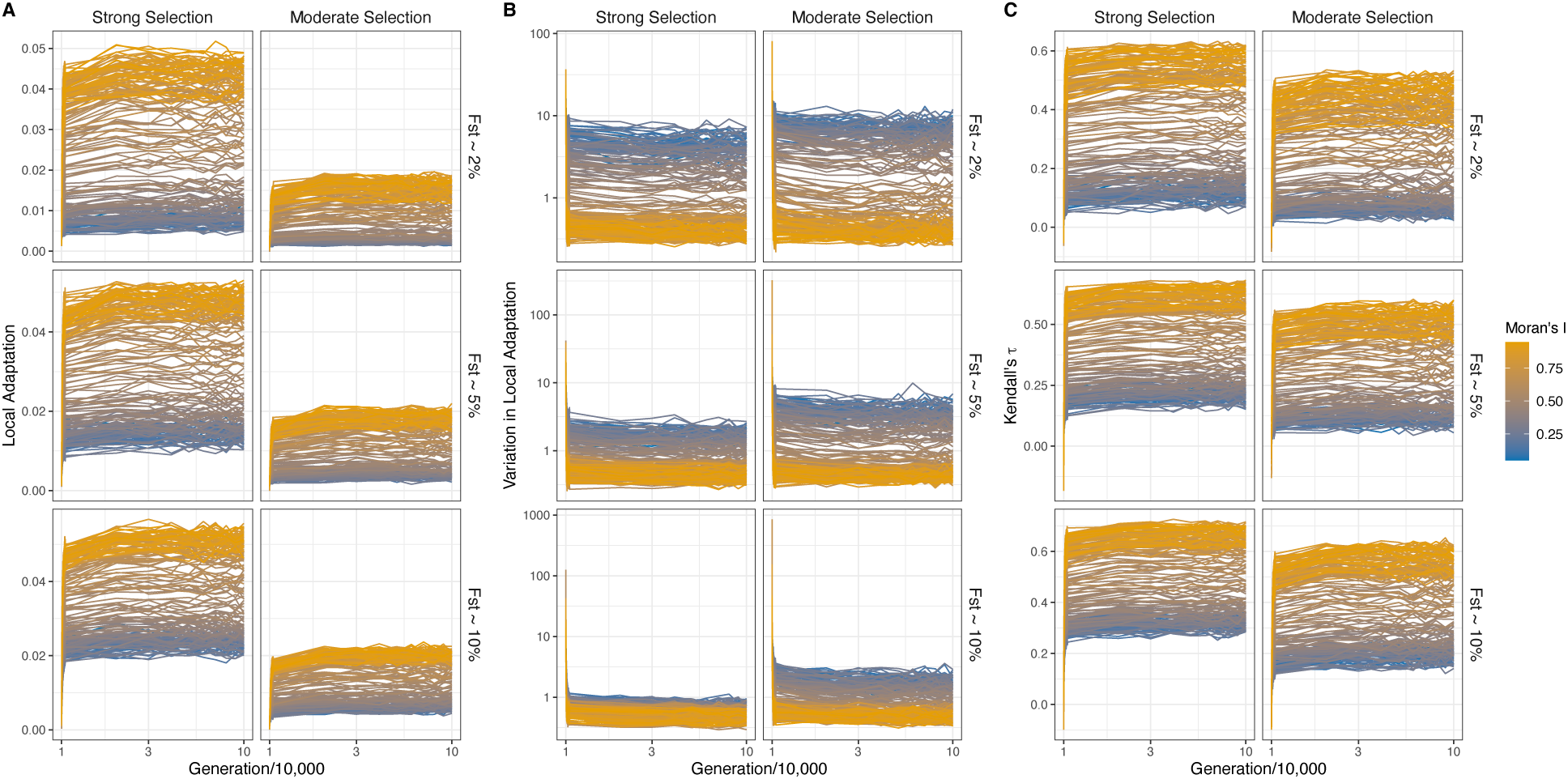
Establishment of local adaptation in the simulations. Panel A) shows the average level of local adaptation across all demes. Panel B) shows the coefficient of variation in local adaptation across demes. Panel C) shows the Kendall’s tau rank correlation between phenotypes and local optima.

**Supplementary Figure 10.**
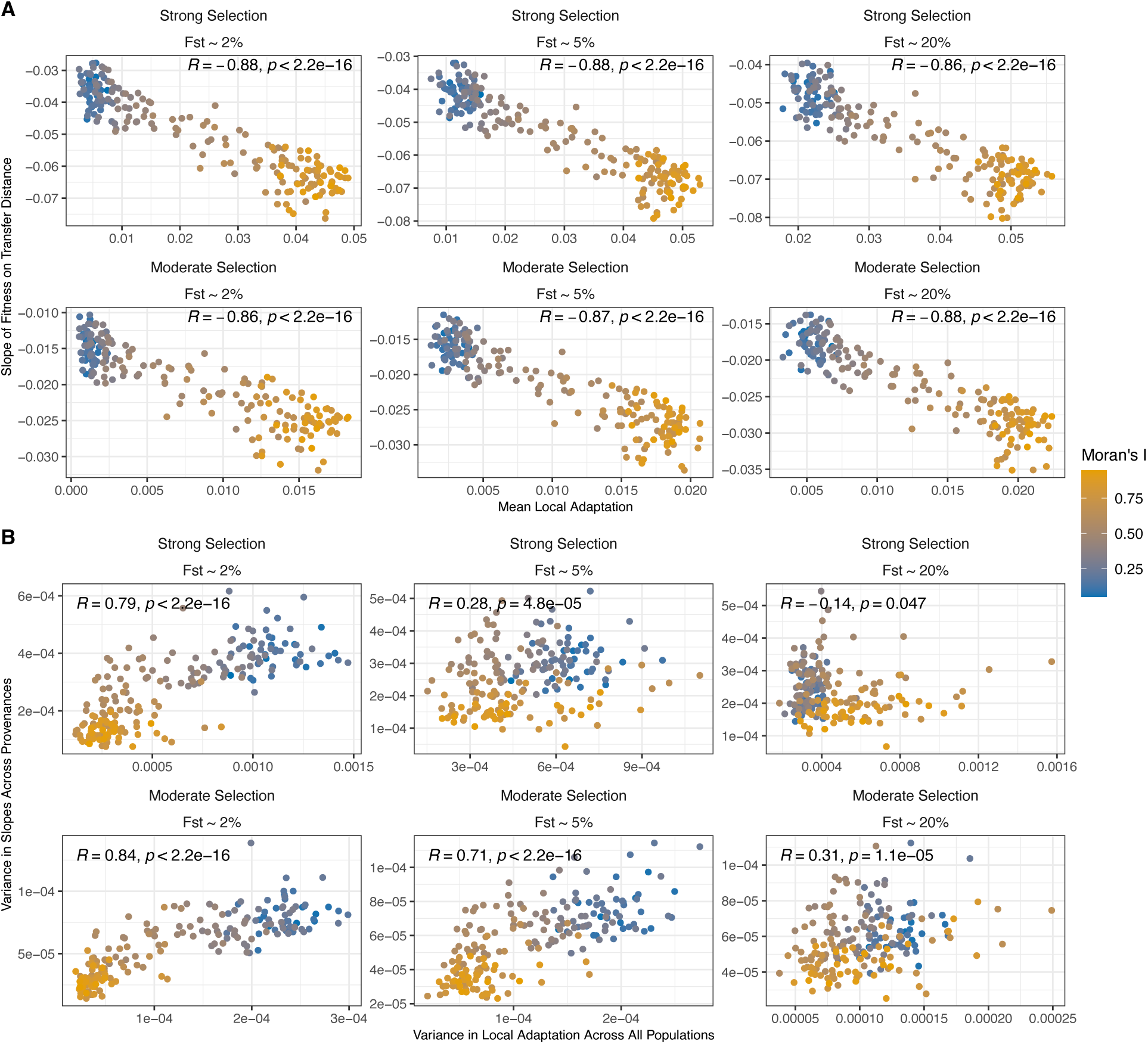
Comparing local adaptation summary statistics to results of linear models applied to simulated provenance trials. Panel A compared the slopes of the relationship between relative fitness and transfer distance in simulated provenance trials to home-versus-away measure of local adaptation described by Blanquart et al., (2013). Panel B compares the variance in slopes across provenances to the variance in local adaptation across all populations. In both panels, each point summarises analyses from a single simulation. The Spearman correlation coefficient and the associated *p*-value are shown within each cell.

**Supplementary Figure 11.**
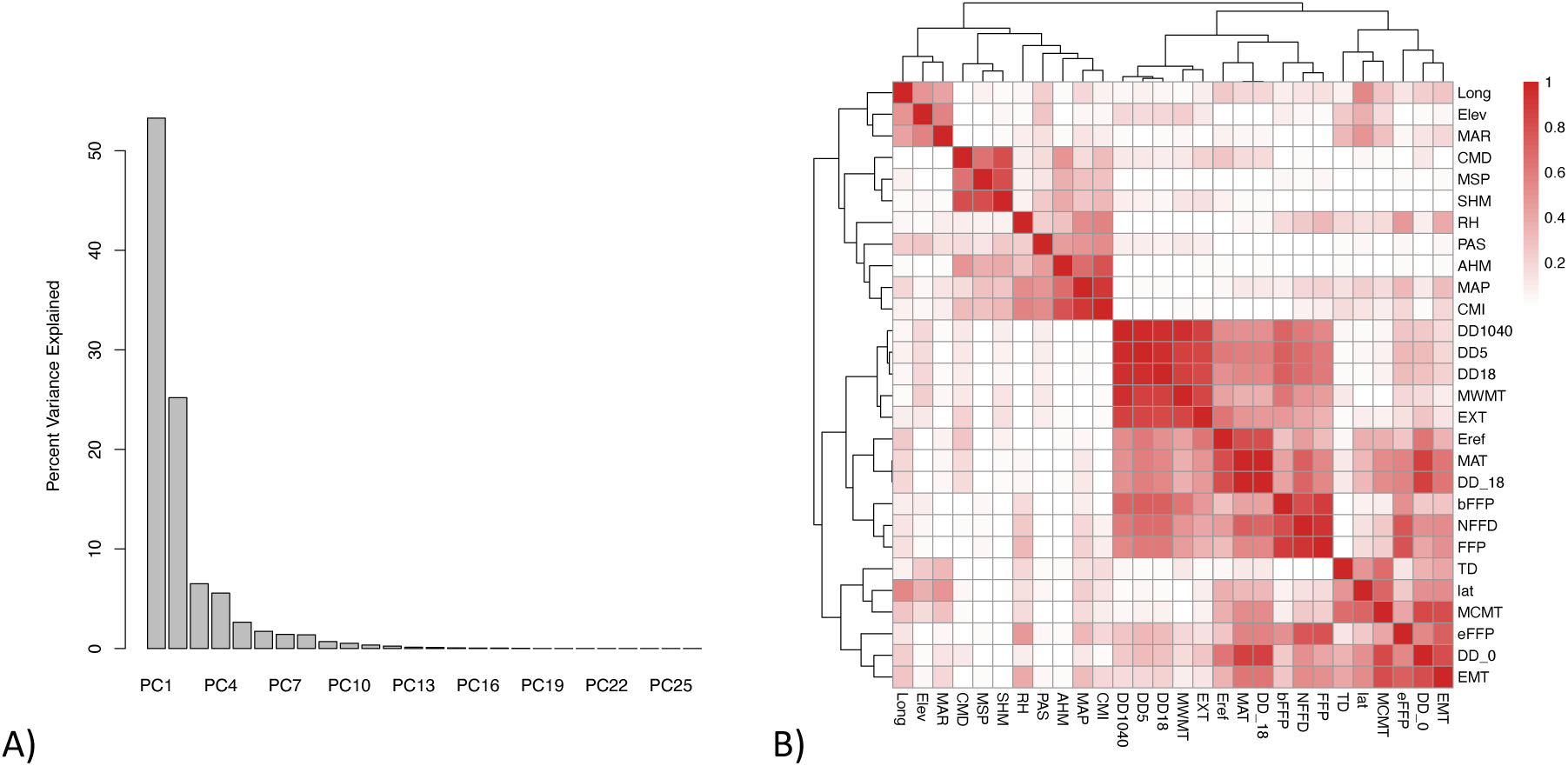
A) Percent variance explained by the principal component analysis conducted on climatic/environmental variation in the Illingworth trial data. B) The correlation matrix for the 28 climatic/environmental variables for planting sites and provenances in the Illingworth trial. A key to the abbreviations for the 25 annual climatic variables from ClimateBC along can be obtained from https://climatebc.ca/Help2. Additionally, latitude (lat), longitude (Long) and elevation (Elev) are included.

**Supplementary Figure 12.**
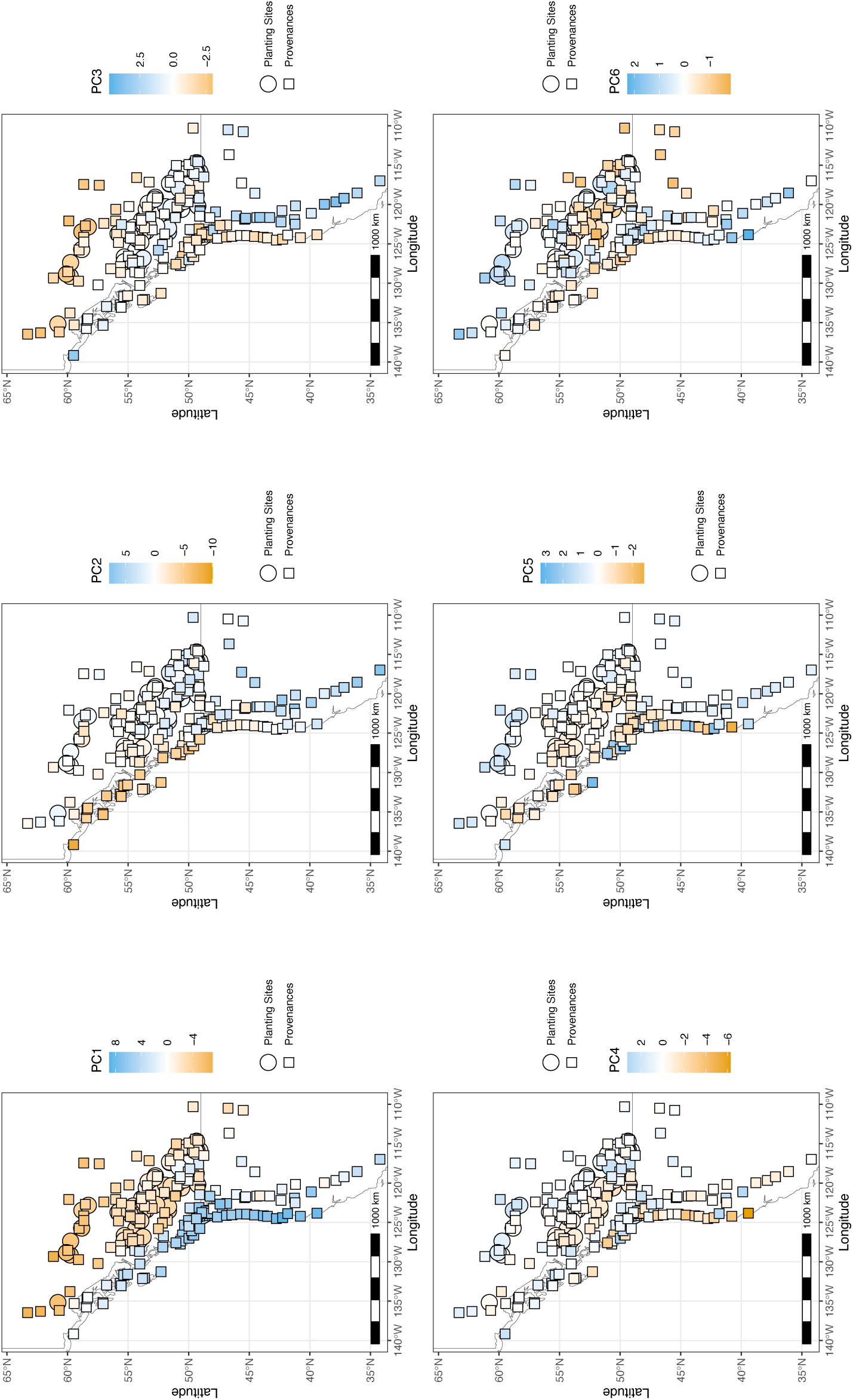
The spatial pattern of loadings onto the first 6 principal components of environmental/climatic variation across provenances and planting sites in the Illingworth Trial. The first 6 principal components explained a total of 95% of the variation in the data.

